# Graph transformer for ancient ancestry inference

**DOI:** 10.64898/2026.04.05.714076

**Authors:** Cole Shanks, David Bonet, Marçal Comajoan Cara, Alexander Ioannidis

**Affiliations:** Genomics Institute, University of California, Santa Cruz, 2300 Delaware Avenue, Santa Cruz, 95060, CA, USA; Department of Electrical Engineering and Computer Science, University of California, Berkeley, 1156 High Street, Berkeley, 94720, CA, USA; Department of Genetics, Stanford Medical School, 291 Campus Drive, Stanford, 94305, CA, USA

## Abstract

Local ancestry inference classifies segments of DNA in admixed individuals by their originating population. However, as the date of admixture becomes older, these segments become shorter and determining their ancestry becomes increasingly difficult. This limits many existing segment-based methods to relatively recent historical admixture events and more highly diverged populations. The rapidly expanding availability of ancient DNA offers a promising opportunity to use these ancient samples as references for local ancestry inference. A recent approach integrates ancient samples into the ancestral recombination graph (ARG) for local ancestry inference. Here, we introduce recent advances in deep learning for graphs into this ARG framework to create ARGMix, a graph transformer that infers local ancestry using the coalescent trees of the inferred ARG. Our approach employs ancient samples as references in the marginal trees to predict local ancestry. We train ARGMix on data reflecting the well-understood ancient European demography and demonstrate improved accuracy and robustness even under demographic misspecification. We then apply ARGMix to an ARG of ancient and present-day European samples for ancestry-specific analyses, finding evidence of continuity between Ötzi the Iceman and present-day individuals from nearby regions.

## 1 Introduction

Local ancestry inference is a critical tool in population genetics [1]. These methods classify the ancestry of DNA segments (local genomic ancestry), enabling analyses in admixed individuals to focus only on those segments from a particular ancestral background (ancestry-specific analyses). This approach is popular for several applications, including increasing power in genome-wide association studies, characterizing ancestry-specific allele frequencies, and reconstructing separate demographic histories for each ancestry [2–4]. Several methods have been developed for local ancestry inference, many of which use hidden Markov models (HMM), and in particular the Li-Stephens HMM [5–7]. Other approaches include Gnomix, which offers a suite of machine learning methods to train a base layer and an additional smoothing layer to improve accuracy [8]. A recent development in population genomics is the inference of ancestral recombination graphs (ARGs) that scale to thousands of samples [9, 10]. Using the inferred ARG, AncestralPaths trains a neural network on the genealogical nearest neighbors with a simulated European demography as ground-truth. This approach utilizes ancient DNA (aDNA) from different time depths as references for ancestry [11]. In this framework, local ancestry is interpreted as a path of successive coalescent events, yielding higher accuracy for older admixture times compared to conventional sequence-based approaches. Importantly, this approach enables the usage of admixed individuals that are informative of their respective ancestries, increasing the size of the reference panel dramatically for some applications. This enables accurate local ancestry inference for the ancient ancestries that formed West Eurasia in present-day individuals, previously infeasible with other methods. This in turn permits the stratification of selection tests by ancestry, mitigating confounding admixture and revealing ancestry-specific signals [11–13]. However, this approach does not model the structure of coalescent trees directly; instead, it linearizes the coalescent events into proportions of genealogical nearest neighbors by population.

Important recent developments in deep learning for graphs include graph neural networks [14, 15], and, still more recently, graph transformers [16]. Graph transformers extend transformers to graph-structured data, allowing self-attention to act between nodes, capturing long-range dependencies [17, 18]. Graph transformers can perform node classification, graph classification, and graph regression tasks, making their application to ancestral recombination graphs natural. Here we introduce ARGMix, a transformer based on the graph relative positional encoding (GRPE) approach [19]. We define relative position between nodes in the ARG using the time-to-most-recent-common-ancestor (TMRCA) and incorporate node population labels to inform local ancestry. We benchmark ARGMix against AncestralPaths and demonstrate an significant improvement in accuracy and robustness under demographic misspecification. As an application, we show that Ötzi the Iceman clusters most closely with modern-day Bergamo Italians when conditioned on Anatolian ancestry, demonstrating continuity of his ancestry with present-day individuals from nearby regions [20]. Additionally, we revisit selection of HLA-DRB1*15:01 using CLUES2 [21], which is associated with high risk of multiple sclerosis and underwent positive selection in the past, but has more recently undergone strong negative selection [12, 22].

## 2 Results

### 2.1 A Graph Transformer for local ancestry inference

ARGMix adapts GRPE to apply a transformer to marginal coalescent trees of the ancestral recombination graph [19]. In the original GRPE formulation, the model included edge encodings in addition to distances between nodes. In this application, a natural way to represent the topological relationships between haplotypes is the TMRCA, so our approach encodes only node-to-node relationships. This model uses the coalescent times (in generations) to relate tokens and modulate attention, wherein each token represents a haplotype in the marginal tree. This design also allows nodes to carry any relevant features. For local ancestry inference, nodes are tokenized with their population labels to inform ancestry. The resulting model focuses on ancestral topology and divergence between haplotypes, rather than local edge connectivity. Further details on how the TMRCA is incorporated into the positional encoding are given in the Methods (Section 4).

Graph transformers can facilitate several graph-related tasks. In the context of local ancestry, one approach might be to perform node classification by applying a transformer to the entire coalescent tree, but this does not scale with standard self-attention (*O*(*n*^2^) in time) [17]. Instead, we turned the task of predicting local ancestry into a subgraph classification problem. This sub-sampling approach is similar to existing graph neural network methods that scale to large graphs, improving scalability [23]. We extracted subgraphs containing specified references informative of ancestry (excluding other extant haplotypes that we may be predicting) and inferred ancestral haplotypes (internal nodes) around the extant haplotype (leaf node) being classified (Figure 1A). Specifically, we selected the nearest reference samples by TMRCA, regardless of ancestry, and a representative set of ancestral haplotypes encoding coalescent events. We considered the four-way demography that created present-day Europe, where the ancestral populations are Anatolian (early farmers), Western, Eastern, and Caucasus hunter-gatherers [24]. In this demographic model of Europe, Anatolian farmers expanded into Europe and eventually admixed with Western hunter-gatherers (WHG), forming an admixed Neolithic farmer population. Later, the Caucasus hunter-gatherers (CHG) admixed with Eastern hunter-gatherers (EHG) just north of the Caucasus, forming the Yamnaya steppe population. Critically, Yamnaya and Neolithic farmer samples are informative of their two respective admixed ancestries and are used as additional references. For clarity, Neolithic farmers here refer to individuals with WHG and Anatolian farmer ancestry.

**Fig. 1:**
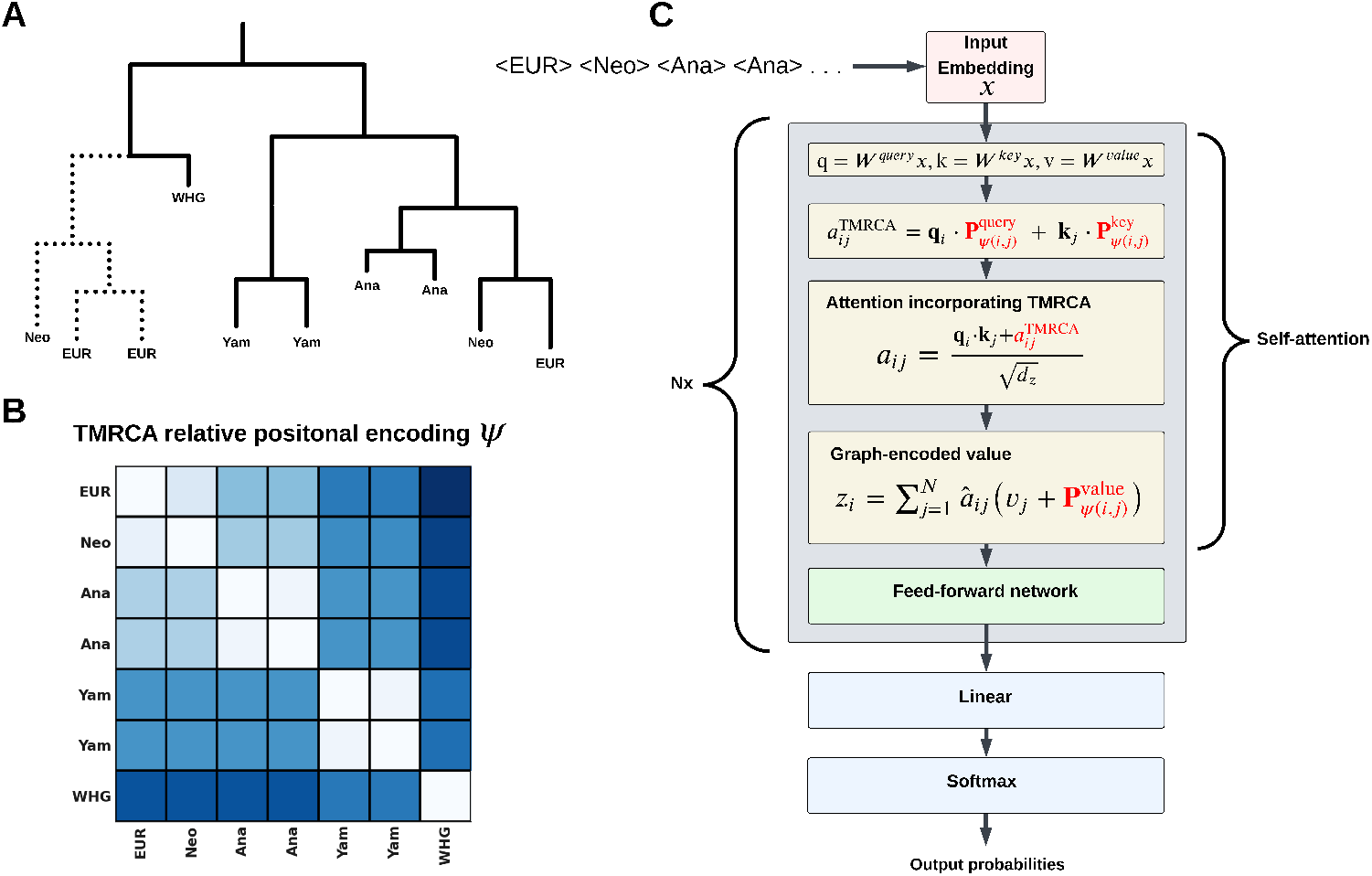
Schematic of ARGMix. (A) Example of a marginal tree inferred with Relate, showing coalescent events among extant haplotypes with population labels. The selected subgraph is indicated with solid lines. (B) TMRCA-based relative positional encoding derived from Relate-inferred marginal trees. (C) ARGMix model architecture incorporating the TMRCA-based relative positional encoding; red highlights indicate modifications that encode TMRCA information.

### 2.2 Training and benchmarks with AncestralPaths

We trained both models on a simulated demography from Irving-Pease et al. 2024 containing 500,000 examples (subgraphs), generated from 5 replicates [11]. The models were trained on the marginal coalescent trees inferred from the simulated data by Relate [9]. We benchmarked our method with AncestralPaths, which also predicts the local ancestry of Neolithic farmers and Yamnaya, and evaluated both on the same samples. We generated additional replicates to assess accuracy, and the largest improvement of ARGMix over AncestralPaths was in present-day Europeans, who carry all four ancestries, with an average gain of 8.23% (Table 1). To show that ARGmix does not overfit to a specific demography and does not underperform in application, we also created misspecified scenarios by randomly perturbing parameters, including population divergence, admixture dates (shifted older), and effective population size (varied by 1-50%). ARGMix demonstrated increased robustness, retaining on average 5.33% higher accuracy than AncestralPaths in present-day individuals (Table 2).

**Table 1:**
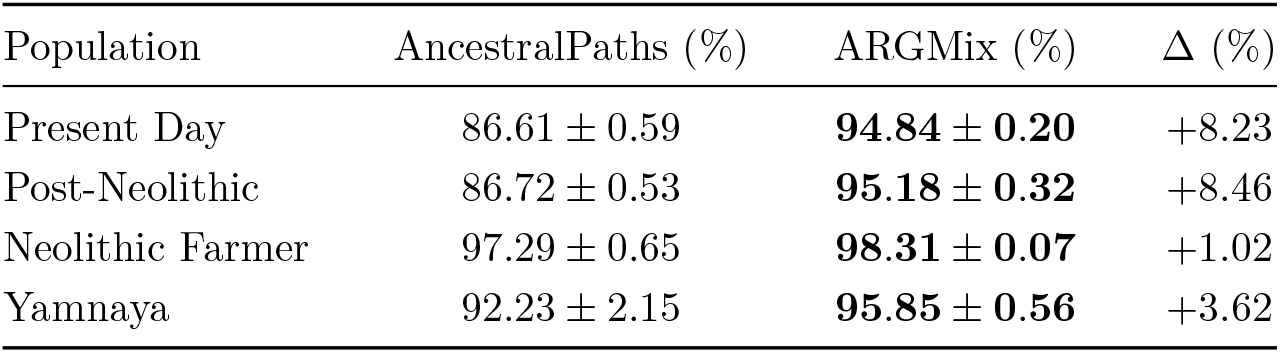
Population-specific accuracy improvement.

**Table 2:**
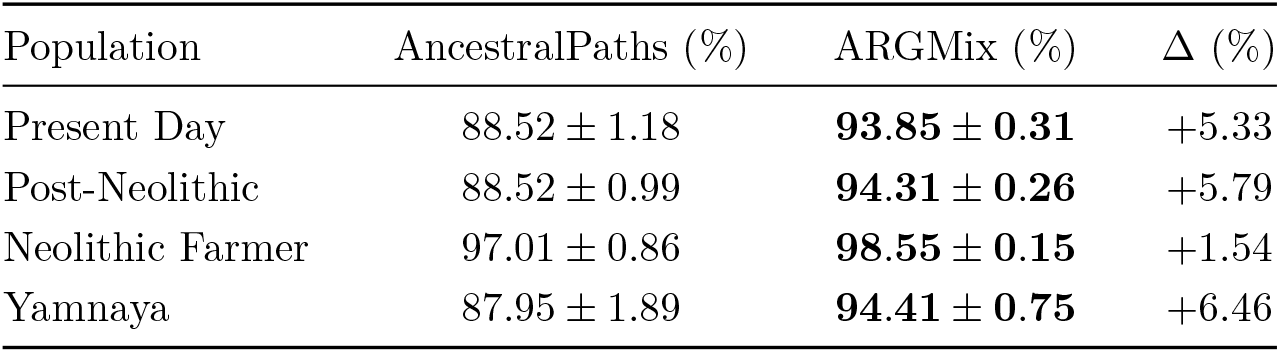
Robustness to misspecified demography.

### 2.3 Anatolian-specific population structure of the Iceman and present-day Europeans

The accuracy of ARGMix allows us to answer ancestry-specific demographic questions. In particular we demonstrate the effectiveness of ARGMix on real data, focusing on the ancient DNA dataset from Allentoft et al. 2024 [24], merged with an additional genome of Ötzi the Iceman [20]. One application of local ancestry inference is the ability to mask-out other ancestries, meaning that the covariance of genotypes between samples can be computed within a specified ancestry only. The Iceman has been reported to be closely related to Sardinians [25, 26]. In a PCA with present-day European populations, the Iceman distinctively clusters with the Sardinians (Figure 2A). However, this pattern likely reflects differences in later admixture proportions–the isolation of Sardinia preserved higher Neolithic farmer ancestry, and the Iceman can be understood as an early Neolithic farmer (Supplementary Figure 5). The question of which European populations are the most closely related descendants to Ötzi the Iceman, that is share the most similar genetics in their relevant Neolithic farmer DNA segments, remains open. Our method allows us to now answer this question. We mask all genotypes (set haplotypes of other ancestries to missing in the VCF) except Anatolian ancestry and then performed ancestry-specific PCA, which accommodates for missingness and controls for differences in admixture proportions. The Iceman now clusters close to modern day Bergamo Italians, that is, the present-day population geographically closest to the site in the Ötztal Alps where the Iceman was found (Figure 2B). We verified this finding by using TreeMix on the Anatolian-specific allele counts we are able to recover using our method [27]. Again Bergamo Italians and the Iceman form the closest clade (Supplementary Figure 4). This suggests genetic continuity in the northern Italian region since the time of the Neolithic farmers and demonstrates the importance of ancestry-specific methods for addressing later confounding admixture when investigating such historical questions.

**Fig. 2:**
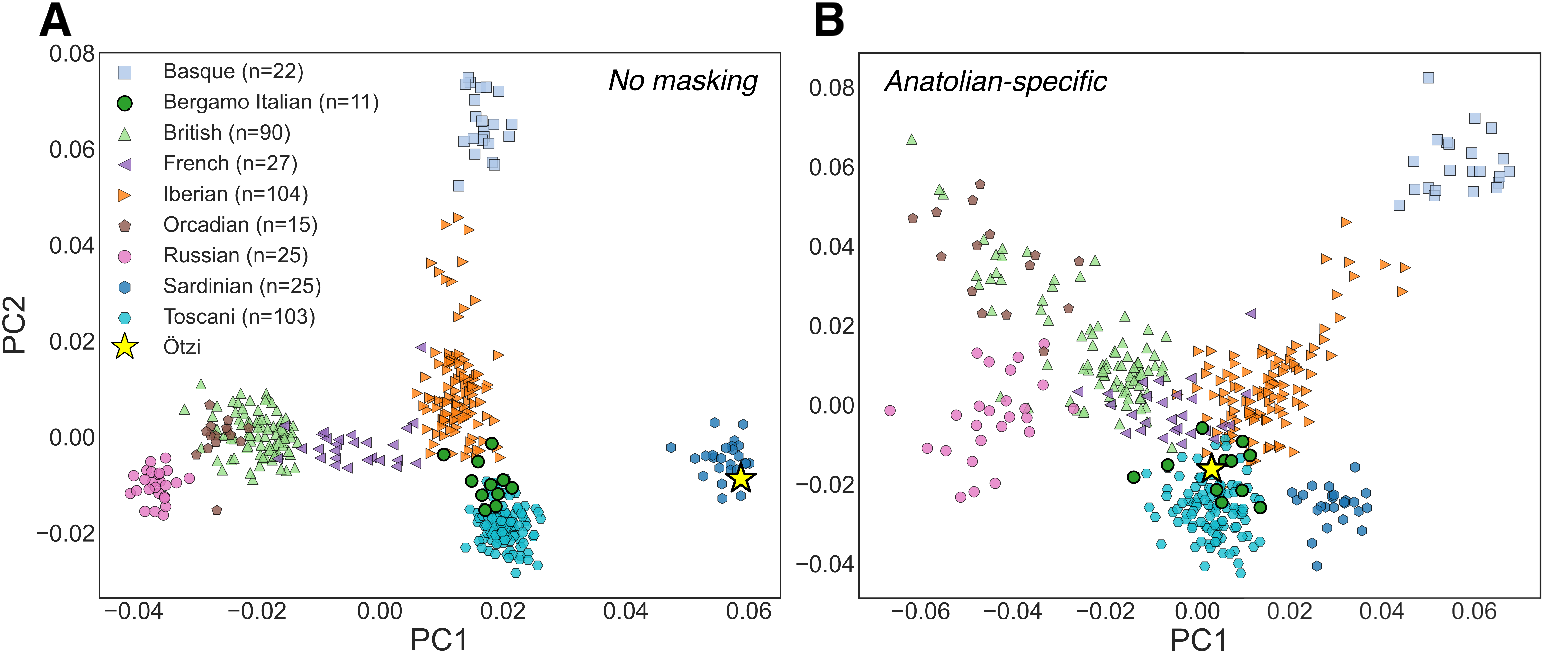
Population structure of present-day Europeans and the Iceman. (A) Principal component analysis (PCA) of the Iceman and European populations from the Human Genome Diversity Project and the 1000 Genomes Project. (B) Anatolian-specific PCA of the same populations generated by masking non-Anatolian ancestry.

To further validate ARGMix, we computed f_4_ statistics to capture admixture in Italy from the Iron Age to present-day Toscani individuals [28] [29], but here again using an ancestry specific approach (Supplementary Figure 6). We tested admixture with Bronze Age Levantine samples and reproduce the known shifts in Italy with increased Mediterranean and Middle East ancestry during the Imperial Roman era due to the large population centers in the Levant and Asia Minor [29, 30]. We evaluated statistics before and after masking for Anatolian ancestry for comparison. Notably, increased European ancestry reduced the Anatolian-like ancestry, as reflected in the unmasked statistic f_4_(Toscani, Italy Iron Age; Bronze Age Levant, Italy Neolithic), which yielded *Z* = −5.6. However, when all populations except Bronze Age Levant are masked for Anatolian ancestry, the direction reverses (now *Z* = 6.14), consistent with increased allele sharing between the Levant and Toscani after controlling for reduced Anatolian-like ancestry in present-day Italians.

### 2.4 Selection of HLA-DRB1*15:01

AncestralPaths was applied to stratify selection tests by local ancestry, originally using CLUES, which leverages the ancestral recombination graph [11, 12]. One notable allele is rs3135388, which tags HLA-DRB1*15:01 and has undergone strong selection originating in the Caucasus and Yamnaya ancestries [12, 31]. We revisited this allele using the improved CLUES2, which can fit multiple selection coefficients over specified time windows [21]. We used CLUES2 since it is based on ancestral recombination graphs, and can incorporate ancient DNA samples. Following Vaughn et al. 2024 [21], we applied CLUES2 to aDNA with ancestry stratification (see Methods) and used two epochs, generations 0 to 50 and 50 to 400, identifying a negative selection coefficient in recent times (Figure 3). ARGMix infers an allele frequency of 0% for this variant in Anatolian and WHG ancestry, whereas AncestralPaths predicted selection occurring later in WHG ancestry; this discrepancy may reflect misclassified local ancestry in AncestralPaths [12]. These results reinforce the conclusions of [12] in that the positive selection appears to begin close to the formation and expansion of the Yamnaya population. This demonstrates the importance of the additional accuracy of ARGMix in another example of ancestry-specific analyses, in this case differential selection histories in haplotypes deriving from different ancestries.

**Fig. 3:**
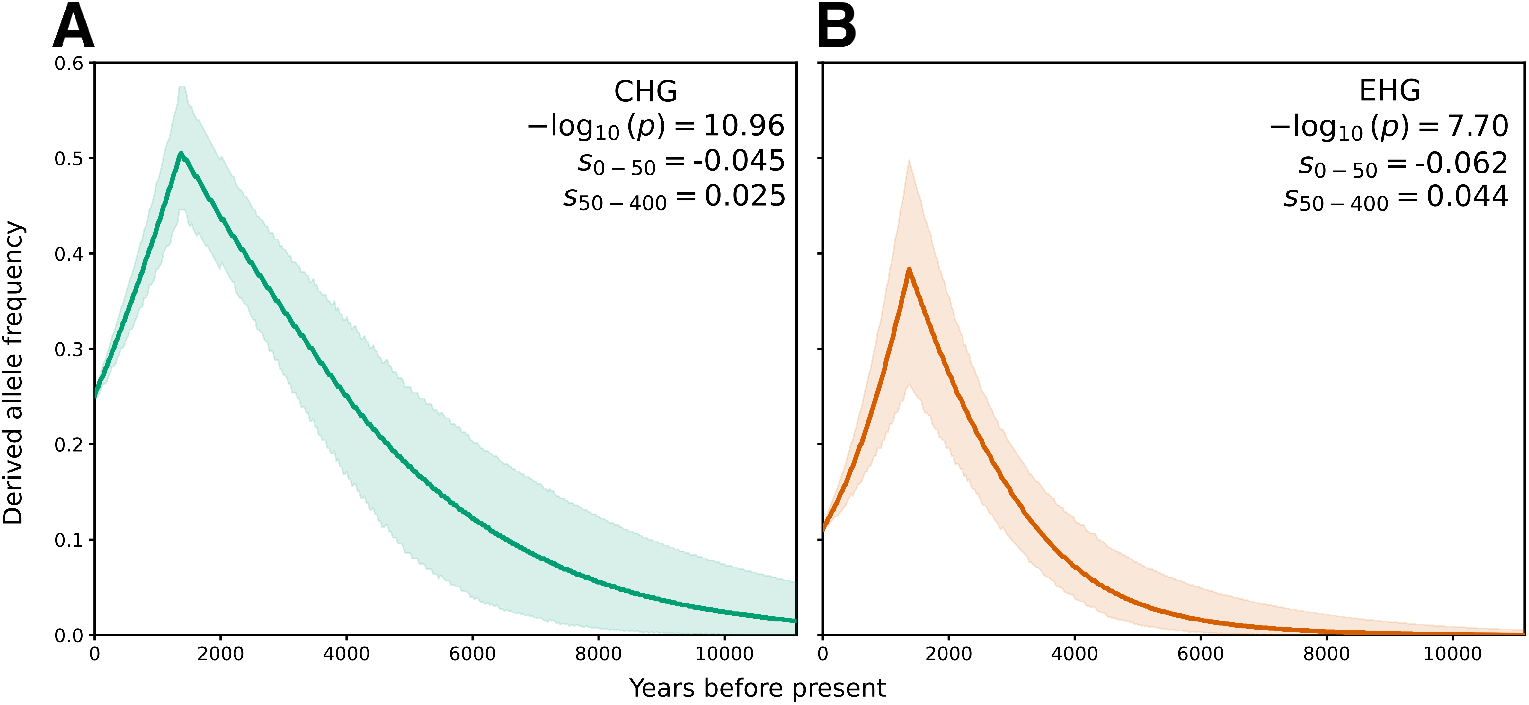
Ancestry-stratified selection test with CLUES2 of rs3135388 tagging HLA-DRB1*15:01. (A) Derived allele frequency through time in CHG ancestry. (B) Derived allele frequency through time in EHG ancestry. Two selection epochs were used, set at generation times 0-50, 50-400. The allele frequency of this variant is predicted to be 0% in Anatolian and WHG ancestries by ARGMix.

## 3 Discussion

Here we introduced ARGMix, a graph transformer for local ancestry inference. By using a richer representation of marginal coalescent trees from the inferred ARG, we achieve improved accuracy over AncestralPaths, especially under demographic misspecification. We show how this method can be applied to ancient samples to demonstrate ancestry-specific population structure. The clustering of the Iceman with modern day Bergamo Italians in their Neolithic farmer DNA segments suggests continuity to the present day in the Alpine region, when combined with the evidence that the Iceman was genetically typical of Tyrol at that time [32]. Additionally, we show that the apparent closer global ancestry relatedness of the Iceman to modern day Sardinians simply reflects later differential admixture proportions, wherein continued admixture reduced, but did not replace, overall Neolithic farmer ancestry in continental Europe compared to the island of Sardinia. Nonetheless, the subtype of Neolithic farmer ancestry present in the Iceman is most similar to the Neolithic ancestry present in nearby modern northern Italians, and not to that found in Sardinians.

We also captured the major transitions in ancestry that affected Italy, made apparent by isolating the Anatolian component. The continued admixture from more northern European populations such as the Lombards, likely reduced the overall proportion of Anatolian ancestry [29], masking the effect of admixture with Middle Eastern ancestry in present-day Toscani in global ancestry based approaches like ADMIXTURE [33]. Here we again show the approach of isolating ancestral segments and analyzing them separately can identify notable admixture that has been confounded by later events, especially those substantially changing the overall global ancestry proportions of individuals.

Local ancestry methods can de-convolve the frequency of a genetic variant in the DNA segments of different origins, which is particularly useful for analyzing selection when it has a different history in the different ancestries. This is important to understand the history of genetic disease and provides insight into sources of selective pressures. We applied ARGMix in the context of selection in different ancestral segments, identifying strong and recent negative selection for HLA-DRB1*15:01, a strong risk factor for multiple-sclerosis [22]. This is consistent with previous work that identified its early origin in the Caucasus, followed by continued positive selection until the last 2000 years, when negative selection was observed [12, 31]. This positive selection continued, possibly reflecting a different pathogen environment with the arrival of steppe ancestry, which rapidly changed again in the recent past. [31] find that the selection of this variant is stronger in the north relative to the south of Europe. [31] report the estimated allele frequencies through time under a 4-way admixture model using qpAdm. Under the 4-way model similar to the one used here, the frequency of HLA-DRB1*15:01 continues to increase until 500-2500 years before present (bp) in EHG ancestry (40% frequency), but rapidly declines in CHG ancestry (9% frequency in 500-2500 bp). Here, using ARGMix and CLUES2 we show a decline in the allele frequency in both EHG and CHG ancestry. This reflects the advantage of using an accurate local ancestry method for such a task. Although we do not explore it here, the more robust local ancestry inference from ARGMix could potentially also identify selection missed by AncestralPaths.

Effectively applying deep learning has remained a challenge in population genetics. A fundamental issue is the lack of ground truth and the reliance on simulated data. Our TMRCA-based relative positional encoding could be expanded into a broader framework for deep learning on ancestral recombination graphs. A notable method is SIA, which uses an LSTM on the number of lineages extracted from the ARG to identify selection [34]. This method outperformed CLUES, which also utilizes the ARG [35]. Similarly, the graph transformer framework described here could be applied to selection with few modifications, such as replacing the population label with the derived-allele state and performing graph regression instead. Here we used the standard self-attention mechanism, which can have a prohibitive runtime for longer sequences. Many methods address this in natural language processing, but we presume that a tree-specific approach may be most effective [36]. We think this graph transformer approach is promising and could be extended to several applications in population genomics given sufficient modifications and development. Additionally, a similar framework could be applied to massive phylogenetic trees beyond ARGs [37, 38].

Data augmentation is common in machine learning, and we applied it only lightly here [39]. Creating training data to better mimic real data is a point of future work that could greatly increase accuracy. Several approaches address these issues besides augmentation, including domain adaptation, test-time adaptation, and fine-tuning [40–42]. We did not explore these here, but the viability of domain adaptation has been demonstrated in population genomics with SIA [43] and PopGenAdapt [44]. Approaches that can learn from real data and address the lack of ground truth in population genomics are promising directions, instead of relying solely on simulations that may never fully capture covariate shifts and unknown biases [45].

## 4 Methods

### 4.1 Graph transformer using TMRCA relative positional encoding

For each sample *i* in a tree, we select the nearest reference haplotypes by time-to-most-recent-common-ancestor (TMRCA), excluding post-Neolithic and present-day European samples and the three immediate ancestral haplotypes. Population labels are tokenized for transformer inputs. This model is based on the graph relative positional encoding introduced by Park et al. 2022 with minor modifications in which we drop encodings related to edges [19]. For self-attention, the linear projections are given by

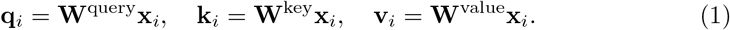

We then define the clamped, asymmetric relative TMRCA

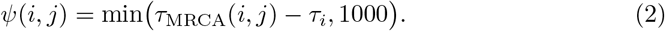

We define an encoding for query, key, and value to represent the relationship between haplotypes by TMRCA,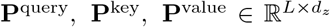. Then an attention map is defined by relative TMRCA that incorporates haplotype features or population labels,

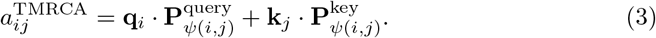

Then the attention map is added to the scaled dot product,

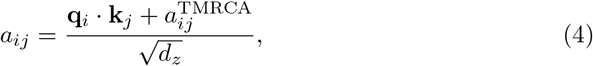

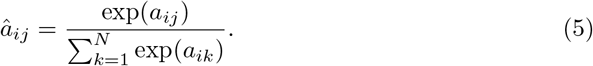

We retain the graph-encoded values and similarly drop any edge information:

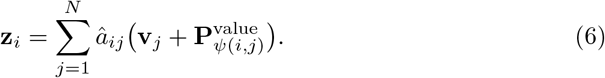

Then **z**_*i*_ is passed to the feedforward neural network. The remainder of the model uses multi-head self-attention.

### 4.2 Simulated data generation

We generated training data using the same simulation as Irving-Pease et al. 2024 [11]. We modified the msprime simulations to exclude Bronze Age Anatolians [46], which were not relevant to our dataset. Similarly, we increased the number of simulated samples to reflect the real dataset we constructed. To better mimic imputation error in ancient DNA, we added a genotype error of 0.001 across all samples, including present-day samples [47]. We generated misspecified demographic scenarios by perturbing the original model of present-day Europe, shifting event timings and effective population size. We randomly varied effective population sizes by an increase or decrease of 1-50%. Admixture timings were increased by 1-25% so as not to break the demography relative to sample times, and population divergences were varied by 1-25% in either direction. We generated all of these simulations using the recombination map for chromosome 22. We ran Relate with an effective population size of 30,000 and a mutation rate of 1.25 ×10^*−*8^ on the simulated data and extracted the ground-truth local ancestry using the simulated ancestral recombination graphs.

### 4.3 Training and evaluating AncestralPaths and ARGMix

We reimplemented AncestralPaths in PyTorch [48], using the proportion of genealogical nearest neighbors (GNN) combined with average sample time [11]. We trained both ARGMix and AncestralPaths on the same 500,000 examples over five replicates. To mitigate the redundancy from local correlation between trees we skipped every ten trees. For validation, we used a separate replicate, skipping every 100 trees and using 100,000 examples. For AncestralPaths, we used the default parameterization that aggregates the GNNs up to five nodes from the extant leaf. For ARGMix, we used 25 of the nearest references by TMRCA plus three inferred ancestral nodes, yielding a 29-node subgraph including the extant leaf being classified. For both models we took the best model by validation loss over all the epochs for further evaluation. To initially evaluate the accuracy, we tested on three additional replicates extracting 100,000 samples from each. We then evaluated five misspecified replicates, also with 100,000 samples and no skipped trees.

### 4.4 Imputation of Ötzi the Iceman

We prepared the large MesoNeo aDNA dataset from Allentoft et al. 2024 [24]. First, we filtered down to the 1492 samples that passed the authors’ quality control. We then restricted to Western Eurasian samples already painted by AncestralPaths. Finally, we filtered for variants with an INFO *>* 0.8. To incorporate the Iceman we started with the BAM file available from Wang et al. 2023 [20]. Using GLIMPSE [49], we applied the same imputation parameters as Allentoft et al. 2024 [24]. However, instead of using the 1000 Genomes Project phase 3 alone, we used a merged Human Genome Diversity Project and 1000 Genomes Project reference panel for imputation [50–52]. This provided additional present-day European populations, such as Sardinians, who have retained more Neolithic farmer ancestry [53]. After imputation, we merged present-day European populations and the Iceman on the intersection of well-imputed variants in the MesoNeo dataset. This yielded 3,720,746 variants total and 1611 samples. All the simulations were generated assuming this sample count. To address differences in haplotype phase, we rephased the final variant set with Beagle 5.4 [54].

### 4.5 Application of ARGMix to ancient DNA and to Ötzi

To construct a reference panel for ARGMix and AncestralPaths, we used the references available in the supplement of Irving-Pease et al. 2024 [11]. We further separated the early farmers and Anatolian farmers to identify those suitable as references for Anatolian ancestry alone. Using 255,816 filtered variants (minor allele frequency exceeding 10% and pruned for linkage-disequilibrium), we ran ADMIXTURE with four clusters on the 1611 samples (Supplementary Figure 2) and required the WHG component to be less than 0.03. The remaining farmers were considered Neolithic farmers [33]. Otherwise we used the same samples as references for the remaining ancestries. Notably, we excluded Bronze Age Anatolians from the simulation used by Irving-Pease et al. 2024 [11]. This final reference set determined simulation parameters, namely sample times and per-population sample sizes (Supplementary Figure 1). In total, we had 22 Anatolians, 13 CHG, 42 EHG, 159 Neolithic Farmers (including the Iceman), 48 WHG, 13 Yamnaya. Additionally, we included 654 post-Neolithic samples and 660 present-day Europeans from HGDP and 1KGP [52]. For all ancestry-specific analyses, we used snputils [55], including PCA and *f* -statistics. To apply CLUES2 we followed the approach of Vaughn et al. 2024 for performing the ancestry-stratified selection tests [21]. For rs3135388 we partitioned each of the ancient haplotypes based on the local ancestry inferred by ARGMix. If a sample was not included in local ancestry inference but was used as a reference, we also included its haplotypes. Then, we computed the frequency of the allele in present-day Europeans conditioned on local ancestry. We used the default parameters otherwise.

## Supplementary information

Figs. S1 to S6.

## Acknowledgements

We thank Martin Sikora for helpful feedback on the selection of local ancestry references. This work was supported by the National Human Genome Research Institute T32HG012344 grant.

## Notes

https://github.com/AI-sandbox/argmix

### Competing Interest Statement

The authors have declared no competing interest.

## References

[1] Sun, Q. et al. Opportunities and challenges of local ancestry in genetic association analyses. The American Journal of Human Genetics 112, 727–740 (2025).

[2] Atkinson, E. G. et al. Tractor uses local ancestry to enable the inclusion of admixed individuals in GWAS and to boost power. Nature Genetics 53, 195–204 (2021).

[3] Kore, P. et al. Improved allele frequencies in gnomAD through local ancestry inference. Nature Communications 16, 8734 (2025).

[4] Browning, S. R. et al. Ancestry-specific recent effective population size in the americas. PLOS Genetics 14, 1–22 (2018).

[5] Yang, Y., Durbin, R., Iversen, A. K. N. & Lawson, D. J. Sparse haplotype-based fine-scale local ancestry inference at scale reveals recent selection on immune responses. Nature Communications 16, 2742 (2025).

[6] Browning, S. R., Waples, R. K. & Browning, B. L. Fast, accurate local ancestry inference with FLARE. American Journal of Human Genetics 110, 326–335 (2023).

[7] Li, N. & Stephens, M. Modeling linkage disequilibrium and identifying recombination hotspots using single-nucleotide polymorphism data. Genetics 165, 2213–2233 (2003).

[8] Hilmarsson, H. et al. High Resolution Ancestry Deconvolution for Next Generation Genomic Data (2021). Preprint at https://www.biorxiv.org/content/10.1101/2021.09.19.460980v1.

[9] Speidel, L., Forest, M., Shi, S. & Myers, S. R. A method for genome-wide genealogy estimation for thousands of samples. Nature Genetics 51, 1321–1329 (2019).

[10] Kelleher, J. et al. Inferring whole-genome histories in large population datasets. Nature genetics 51, 1330–1338 (2019).

[11] Irving-Pease, E. K. et al. The selection landscape and genetic legacy of ancient Eurasians. Nature 625, 312–320 (2024).

[12] Barrie, W. et al. Elevated genetic risk for multiple sclerosis emerged in steppe pastoralist populations. Nature 625, 321–328 (2024).

[13] Pandey, D., Harris, M., Garud, N. R. & Narasimhan, V. M. Leveraging ancient DNA to uncover signals of natural selection in Europe lost due to admixture or drift. Nature Communications 15, 9772 (2024).

[14] Wu, Z. et al. A Comprehensive Survey on Graph Neural Networks. IEEE Transactions on Neural Networks and Learning Systems 32, 4–24 (2021).

[15] Zhou, J. et al. Graph Neural Networks: A Review of Methods and Applications (2021). Preprint at https://arxiv.org/abs/1812.08434.

[16] Müller, L., Galkin, M., Morris, C. & Rampášek, L. Attending to Graph Transformers (2024). Preprint at https://arxiv.org/abs/2302.04181.

[17] Vaswani, A. et al. Attention is All you Need, Vol. 30 (2017). Advances in Neural Information Processing Systems.

[18] Bahdanau, D., Cho, K. & Bengio, Y. Neural Machine Translation by Jointly Learning to Align and Translate (2016). Preprint at https://arxiv.org/abs/1409.0473.

[19] Park, W., Chang, W., Lee, D., Kim, J. & Hwang, S.-w. GRPE: Relative Positional Encoding for Graph Transformer (2022). Preprint at https://arxiv.org/abs/2201.12787.

[20] Wang, K. et al. High-coverage genome of the Tyrolean Iceman reveals unusually high Anatolian farmer ancestry. Cell Genomics 3, 100377 (2023).

[21] Vaughn, A. H. & Nielsen, R. Fast and Accurate Estimation of Selection Coefficients and Allele Histories from Ancient and Modern DNA. Molecular Biology and Evolution 41 (2024).

[22] Alcina, A. et al. Multiple Sclerosis Risk Variant HLA-DRB1*1501 Associates with High Expression of DRB1 Gene in Different Human Populations. PLoS ONE 7, e29819 (2012).

[23] Hou, Z. et al. GraphMAE: Self-Supervised Masked Graph Autoencoders. Preprint at http://arxiv.org/abs/2205.10803.

[24] Allentoft, M. E. et al. Population genomics of post-glacial western Eurasia. Nature 625, 301–311 (2024).

[25] Keller, A. et al. New insights into the Tyrolean Iceman’s origin and phenotype as inferred by whole-genome sequencing. Nature Communications 3, 698.

[26] Sikora, M. et al. Population Genomic Analysis of Ancient and Modern Genomes Yields New Insights into the Genetic Ancestry of the Tyrolean Iceman and the Genetic Structure of Europe. PLOS Genetics 10, e1004353.

[27] Pickrell, J. K. & Pritchard, J. K. Inference of Population Splits and Mixtures from Genome-Wide Allele Frequency Data. PLOS Genetics 8, e1002967 (2012).

[28] Patterson, N. et al. Ancient Admixture in Human History. Genetics 192, 1065–1093 (2012).

[29] Antonio, M. L. et al. Ancient Rome: A genetic crossroads of Europe and the Mediterranean. Science 366, 708–714 (2019).

[30] Lazaridis, I. et al. A genetic probe into the ancient and medieval history of Southern Europe and West Asia. Science 377, 940–951 (2022).

[31] Akbari, A. et al. Pervasive findings of directional selection realize the promise of ancient DNA to elucidate human adaptation. bioRxiv 2024.09.14.613021 (2024).

[32] Croze, M. et al. Genomic diversity and structure of prehistoric alpine individuals from the Tyrolean Iceman’s territory. Nature Communications 16, 6431 (2025).

[33] Alexander, D. H. & Lange, K. Enhancements to the ADMIXTURE algorithm for individual ancestry estimation. BMC Bioinformatics 12, 246 (2011).

[34] Hejase, H. A., Mo, Z., Campagna, L. & Siepel, A. A Deep-Learning Approach for Inference of Selective Sweeps from the Ancestral Recombination Graph. Molecular Biology and Evolution 39 (2022).

[35] Stern, A. J., Wilton, P. R. & Nielsen, R. An approximate full-likelihood method for inferring selection and allele frequency trajectories from DNA sequence data. PLOS Genetics 15, e1008384 (2019).

[36] Wang, S., Li, B. Z., Khabsa, M., Fang, H. & Ma, H. Linformer: Self-attention with linear complexity (2020). Preprint at https://arxiv.org/abs/2006.04768.

[37] Turakhia, Y. et al. Ultrafast Sample placement on Existing tRees (UShER) enables real-time phylogenetics for the SARS-CoV-2 pandemic. Nature Genetics 53, 809–816 (2021).

[38] Corbett-Detig, R. A Phylogenetic Method Identifies Candidate Drivers of the Evolution of the SARS-CoV-2 Mutation Spectrum. Molecular Biology and Evolution 42 (2025).

[39] Shorten, C., Khoshgoftaar, T. M. & Furht, B. Text Data Augmentation for Deep Learning. Journal of Big Data 8, 101 (2021).

[40] Ganin, Y. & Lempitsky, V. Unsupervised Domain Adaptation by Backpropagation (2015). Preprint at https://arxiv.org/abs/1409.7495.

[41] Liang, J., He, R. & Tan, T. A Comprehensive Survey on Test-Time Adaptation under Distribution Shifts. International Journal of Computer Vision 133, 31–64 (2025).

[42] Goyal, S., Kumar, A., Garg, S., Kolter, Z. & Raghunathan, A. Finetune like you pretrain: Improved finetuning of zero-shot vision models, 19338–19347 (IEEE, Vancouver, BC, Canada, 2023).

[43] Mo, Z. & Siepel, A. Domain-adaptive neural networks improve supervised machine learning based on simulated population genetic data. PLOS Genetics 19, e1011032 (2023).

[44] Comajoan Cara, M., Mas Montserrat, D. & Ioannidis, A. G. PopGenAdapt: Semi-supervised domain adaptation for genotype-to-phenotype prediction in underrepresented populations (2024). Biocomputing 2024.

[45] Shi, Y., Xu, W. & Hu, P. Out of distribution learning in bioinformatics: Advancements and challenges. Briefings in Bioinformatics 26, bbaf294 (2025).

[46] Baumdicker, F. et al. Efficient ancestry and mutation simulation with msprime 1.0. Genetics 220, iyab229 (2022).

[47] Sousa da Mota, B. et al. Imputation of ancient human genomes. Nature Communications 14, 3660 (2023).

[48] Paszke, A. et al. PyTorch: An Imperative Style, High-Performance Deep Learning Library, Vol. 32 (2019). Advances in Neural Information Processing Systems.

[49] Rubinacci, S., Hofmeister, R. J., Sousa da Mota, B. & Delaneau, O. Imputation of low-coverage sequencing data from 150,119 UK Biobank genomes. Nature Genetics 55, 1088–1090 (2023).

[50] Byrska-Bishop, M. et al. High-coverage whole-genome sequencing of the expanded 1000 Genomes Project cohort including 602 trios. Cell 185, 3426–3440.e19 (2022).

[51] Bergström, A. et al. Insights into human genetic variation and population history from 929 diverse genomes. Science 367, eaay5012 (2020).

[52] Koenig, Z. et al. A harmonized public resource of deeply sequenced diverse human genomes. Genome Research 34, 796–809 (2024).

[53] Marcus, J. H. et al. Genetic history from the Middle Neolithic to present on the Mediterranean island of Sardinia. Nature Communications 11, 939 (2020).

[54] Browning, B. L., Tian, X., Zhou, Y. & Browning, S. R. Fast two-stage phasing of large-scale sequence data. The American Journal of Human Genetics 108, 1880–1890 (2021).

[55] Bonet, D. et al. snputils: A high-performance Python library for genetic variation and population structure. bioRxiv (2026). URL https://www.biorxiv.org/content/10.64898/2026.02.28.708618.

