## Supplementary Figures for "Graph transformer for ancient ancestry inference"

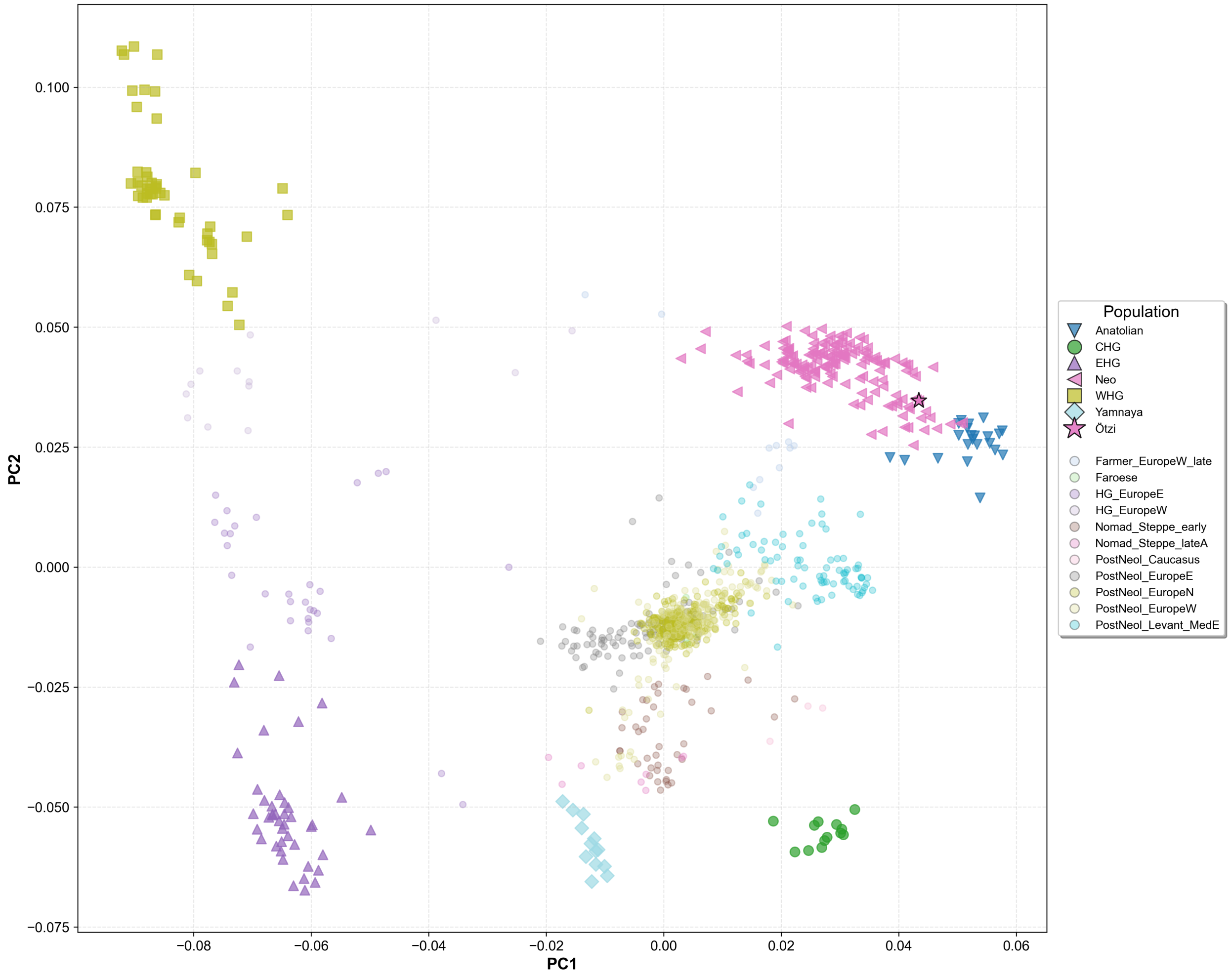


Supplementary Figure S1: PCA of ancient DNA samples. Indicated in bold shapes are references used for local ancestry with ARGMix.

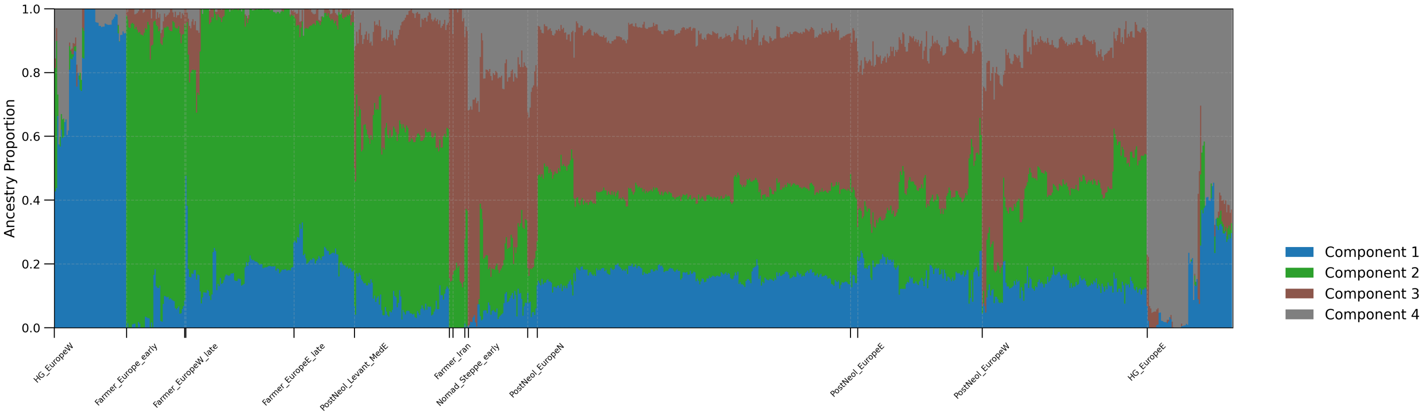


Supplementary Figure S2: Plot of ADMIXTURE unsupervised clustering run on ancient DNA samples with K=4.

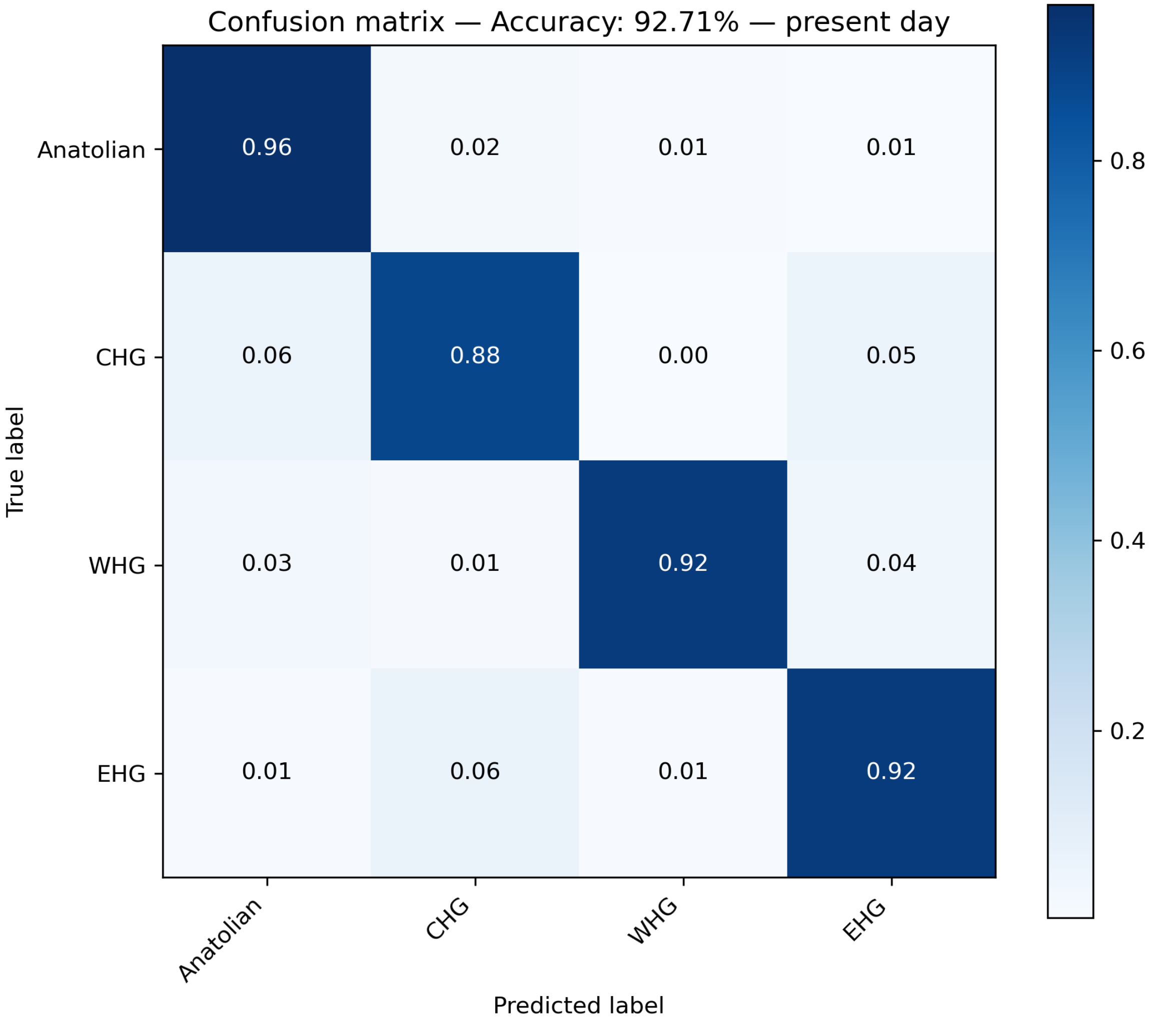


Supplementary Figure S3: Confusion matrix of present-day Europeans for worst-performing misspecified demography nevertheless showing robust accuracy.

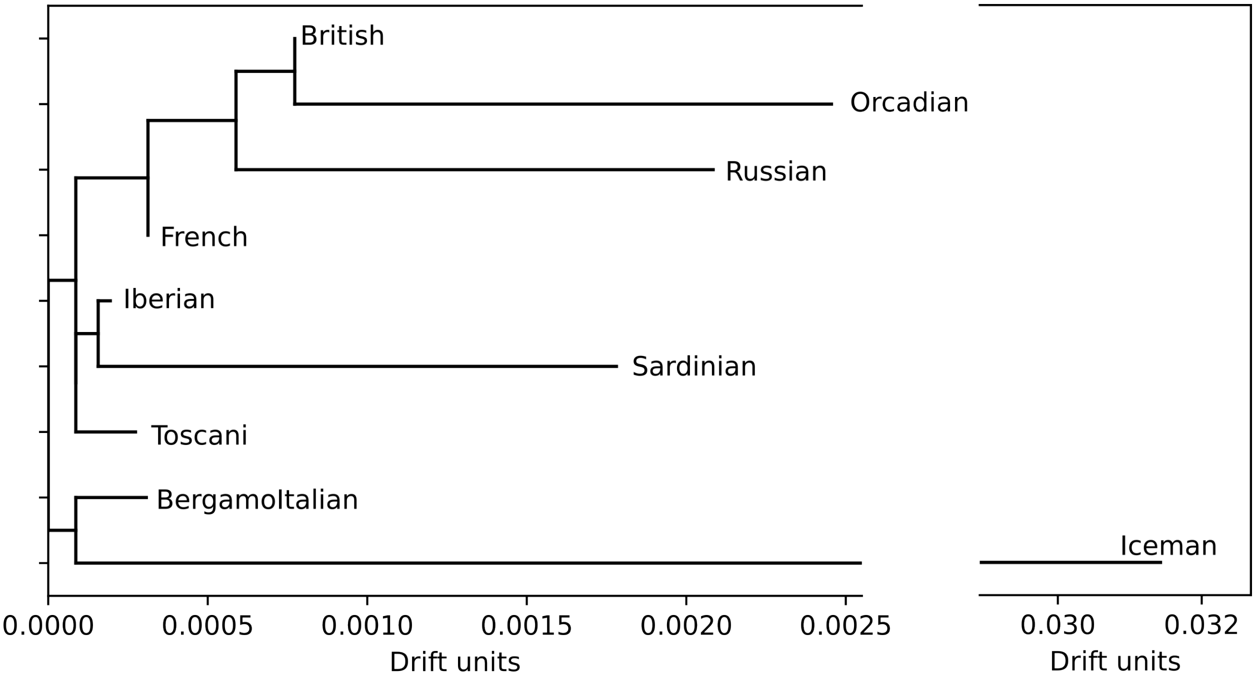


Supplementary Figure S4: Anatolian-specific Treemix results with zero migration edges shows Ötzi the Iceman sharing a clade with modern day northern Italians (Bergamo Italians).

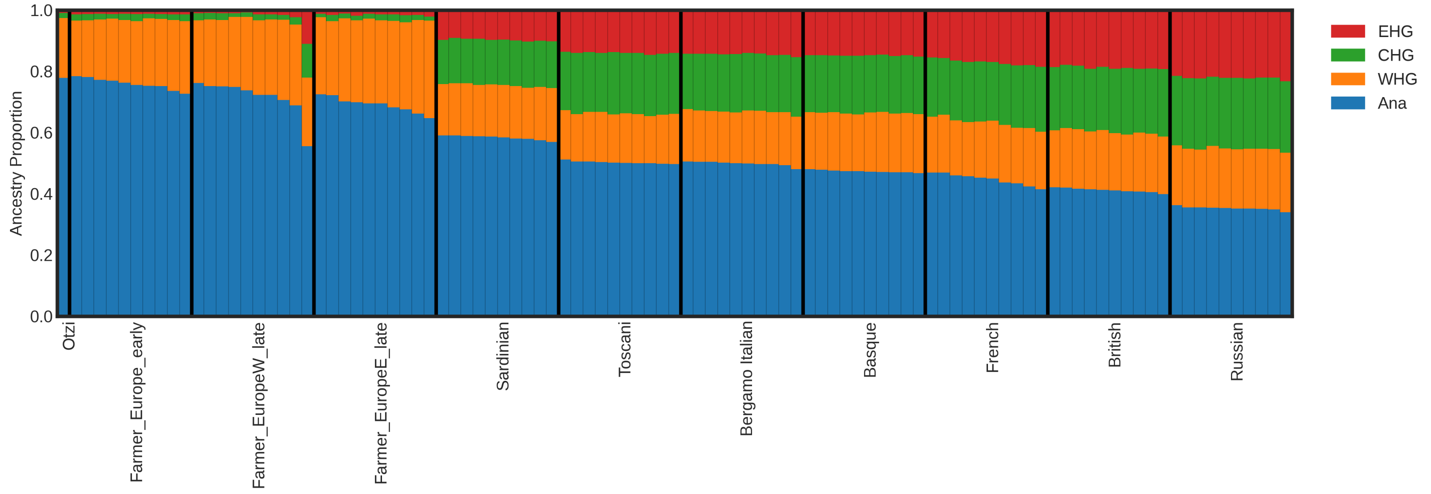


Supplementary Figure S5: Admixture proportions determined by summing of local ancestry calls from ARGMix. Present-day Europeans are included along with early and late Neolithic farmers. Populations are sorted by average Anatolian proportion.

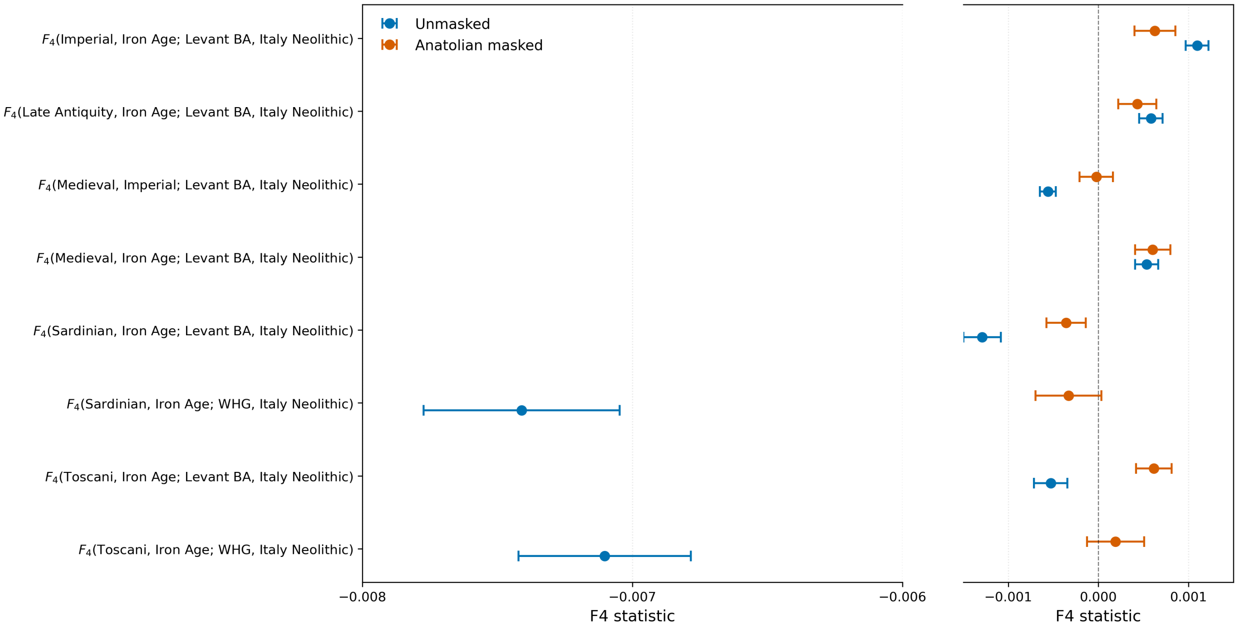


Supplementary Figure S6: Changes in F4 statistics after masking for Anatolian ancestry. Imperial and Iron Age refer to Imperial Italians and Iron Age Italians.
